# A network for self-transcendence derived from patients with brain lesions

**DOI:** 10.64898/2026.03.24.713239

**Authors:** Morgan Healey, Yaser Sanchez-Gama, Mengyuan Ding, James Tanner McMahon, Chase Bourbon, Rumaisa Jesani, Ginger Atwood, Brian Lord, Jay Sanguinetti, Judson Brewer, David Vago, Shan Siddiqi, Franco Fabbro, Cosimo Urgesi, Jared Nielsen, Michael Ferguson

## Abstract

Self-transcendence, the reorientation of experience away from the self toward others, nature, or broader meaning, is a fundamental dimension of human psychology, yet its causal neural architecture remains poorly understood. Here we applied lesion network mapping to 88 neurosurgical patients with pre- and post-operative assessments of trait self-transcendence to identify the distributed brain network whose disruption alters this capacity. The resulting network showed significant spatial correspondence with the default mode network and, at a finer parcellation level, with frontoparietal control subnetworks. Leave-one-out analyses identified posterior midline regions as the most stable correlates of increased self-transcendence following brain lesions. Independent validation against fMRI meta-analyses of self-referential processing, compassion, and ketamine administration, alongside a neuromodulation target previously shown to modulate the sense of self, converged on a consistent model. These findings provide causal evidence for a network architecture in which posterior midline hubs constrain, and brainstem and anterior midline regions facilitate, self-transcendent experience.

## Background

Ego-centric cognition is inherent to human experience and supports adaptive functions related to resource allocation, safety, and goal-directed behavior. At the same time, the capacity to move beyond an ego-bound sense of self, that is, to transcend the self, is a robust feature of human psychology across cultures and contexts (Ge & Yang, 2023; M. Huang & Yang, 2023; Pizarro et al., 2021; Teixeira, 2008). *Self-transcendence* has been conceptualized across a wide range of psychological and philosophical traditions: as a class of prosocial emotional states that reduce focus on oneself and elevate concern for others (Caprara et al., 2012; Pizarro et al., 2021; Stellar et al., 2017; Van Kleef & Lelieveld, 2022), as a stable personality trait (Cloninger et al., 1993), as a transient phenomenological state characterized by ego-dissolution and heightened connectedness (Yaden et al., 2017), and as a spiritual or religious quality (Montemaggi, 2017). Across these perspectives, definitions converge on a common theme: self-transcendence involves a reorientation away from self-focused cognition toward something beyond the self, whether it be other people, the natural world, or a broader sense of meaning and purpose (American Psychological Association, 2023; Garcia-Romeu, 2010; Ge & Yang, 2023; Kitson et al., 2020; Yaden et al., 2017).

While self-transcendence has traditionally been examined through self-report and behavioral approaches, a growing body of work seeks to characterize its neural foundations. Because self-transcendence is defined by a reorientation away from self-focused cognition, it has been theorized as a reconfiguration of self-referential processing at the neural level (Sacchet et al., 2024; Vago & Silbersweig, 2012). Positive psychology has defined the category of self-transcending emotions, some of which have been examined through functional task imaging (Kim et al., 2020). In terms of facilitation in laboratory settings, psychedelics are increasingly being studied in active experimental conditions that induce self-transcendent experience (Carhart-Harris et al., 2012; Watts et al., 2022). The most substantial focus of self-transcendence neuroimaging based research has been on the default mode network (DMN), a large-scale brain system implicated in self-referential thought, autobiographical memory, and internally directed attention (Andrews-Hanna et al., 2010; Buckner et al., 2008). Across meditative and related contemplative states, reduced activity within canonical DMN regions has been reported in association with diminished self-focus and decentering: experiential features central to self-transcendence (Brewer et al., 2011; Garrison et al., 2015; Hanley et al., 2020).

Because functional neuroimaging studies are correlational by design, lesion-based methods provide an important complementary strategy for establishing causal neural substrates of self-transcendence. In a seminal study, Urgesi and colleagues (2010) used voxel-based lesion-symptom mapping in neurosurgical patients assessed before and after brain tumor resection and found that damage to the inferior posterior parietal cortex was associated with changes in trait self-transcendence. This finding provided rare causal evidence linking focal brain disruption to alterations in self-related experiential orientation (Urgesi et al., 2010). This result aligns with broader evidence implicating the inferior posterior parietal cortex, and in particular the temporoparietal junction, in switching between key reference frames: egocentric and allocentric spatial reference, self-other distinction, and theory of mind perspective-taking (Martin et al., 2019, 2020; Orti et al., 2024; Quesque & Brass, 2019).

However, identifying a single anatomical locus may be an overly limited description for a high-order psychological function, such as self-transcendence. The brain’s relationship to higher-order functions is increasingly understood to arise from distributed network interactions rather than isolated region-specific activity (Ferguson et al., 2024; Fox, 2018). Lesion network mapping (LNM) provides a powerful framework for localizing these functions by first identifying cases of focal brain lesions that disrupt the function, then mapping those lesions to a common functional brain circuit (Boes et al., 2015; Fox, 2018). This approach has successfully identified networks underlying a range of complex behaviors, including criminality, general religiosity, and religious fundamentalism (Darby et al., 2018; Ferguson et al., 2022, 2024).

In the present study, we ask: what is the network architecture of self-transcendence and does it correspond to the DMN as predicted by the functional imaging literature? To answer this question we applied LNM to derive a distributed neural network associated with post-surgical changes in self-transcendence, building on the foundational lesion findings of Urgesi et al. (2010).

We then performed a series of analyses to better characterize and validate our derived network. Because of the dichotomy reflected by self-transcendence and self-reference, we compared this network to fMRI meta-analytic maps of self-referential processing. We then triangulated these results with independent lines of external evidence, including fMRI meta-analytic maps of self-transcendent emotions, a fMRI meta-analytic map of ketamine-induced neural activity, and a brain stimulation target implicated to modulate the sense of self.

Given the consistent observation of altered DMN connectivity in self-transcendent states, we tested the hypothesis that self-transcendence-related lesion connectivity would demonstrate spatial association with the DMN. Additionally, we characterized the correspondence of the full network with canonical brain systems using Yeo 7- and 17-network parcellations. Finally, we conducted exploratory comparisons with lesion networks associated with other neurobehavioral conditions. Together, these analyses provide rare causal evidence toward identifying the brain network architecture for self-transcendence.

## Methods

### Deriving the network for self-transcendence

We analyzed a previously published dataset of 88 neurosurgical patients (Urgesi et al., 2010) who completed the Temperament and Character Inventory (TCI) before and after tumor resection (Fig. 1). The primary outcome was pre-to-post-operative change in the TCI Self-Transcendence (ST) subscale, which measures the degree to which an individual identifies with something beyond the self, whether it be nature, humanity, or a higher power (Cloninger, 1994; Cloninger et al., 1993). Details of recruitment, surgical procedures, and psychometric assessment are provided in the original report (Urgesi et al., 2010).

**Figure 1:**
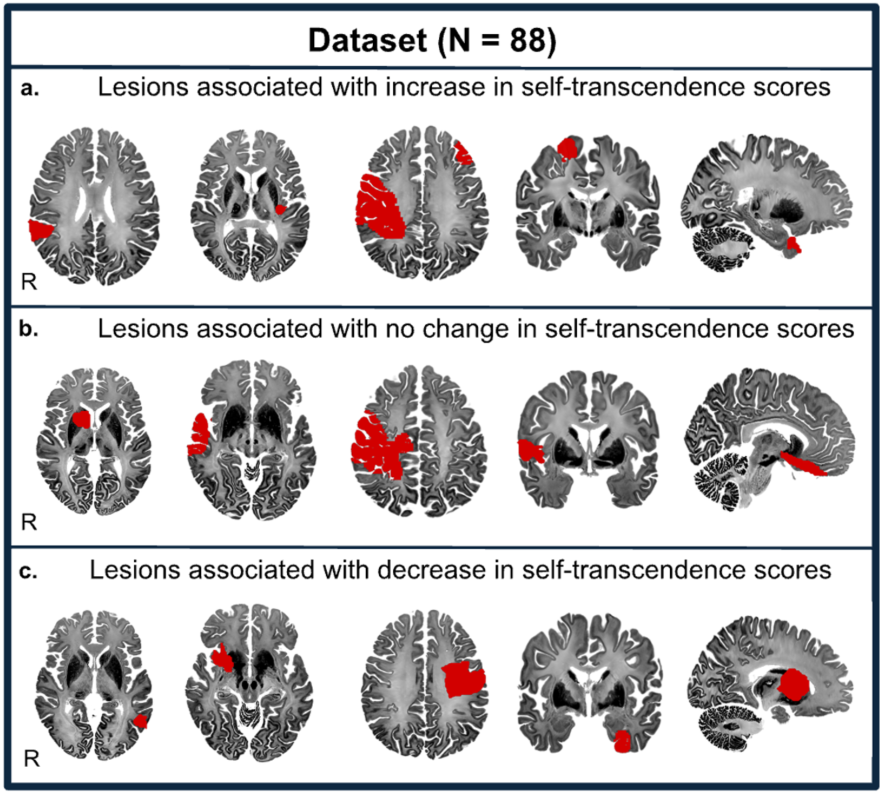
Lesions associated with self-transcendence occur in many different regions. (a) Brain lesions of five patients with the largest increase in self-transcendence scores. (b) Brain lesions of five patients with no change in self-transcendence scores. (c) Brain lesions of five patients with the largest decrease in self-transcendence scores.

Lesion network mapping was performed using established methods (Ferguson et al., 2022; Fox, 2018). This analytic framework links focal lesion data with normative resting-state connectivity to characterize the broader functional architecture underlying complex behavior. Pre-operative structural MRI scans were manually traced to generate binary lesion masks, which were then normalized to MNI152 space using SPM12 with cost-function masking to prevent deformation of lesion tissue. Each mask was visually inspected for accurate registration in both native and standard space. Each lesion mask served as a seed for resting-state functional connectivity analysis computed against a normative connectome dataset (N = 1,000 healthy right-handed participants; 42.7% male; ages 18–35 years; mean age 21.3 years). Voxelwise Pearson correlations between each binarized lesion seed and every brain voxel were calculated and Fisher z-transformed to produce individual lesion-network maps (Fig. 2).

**Figure 2:**
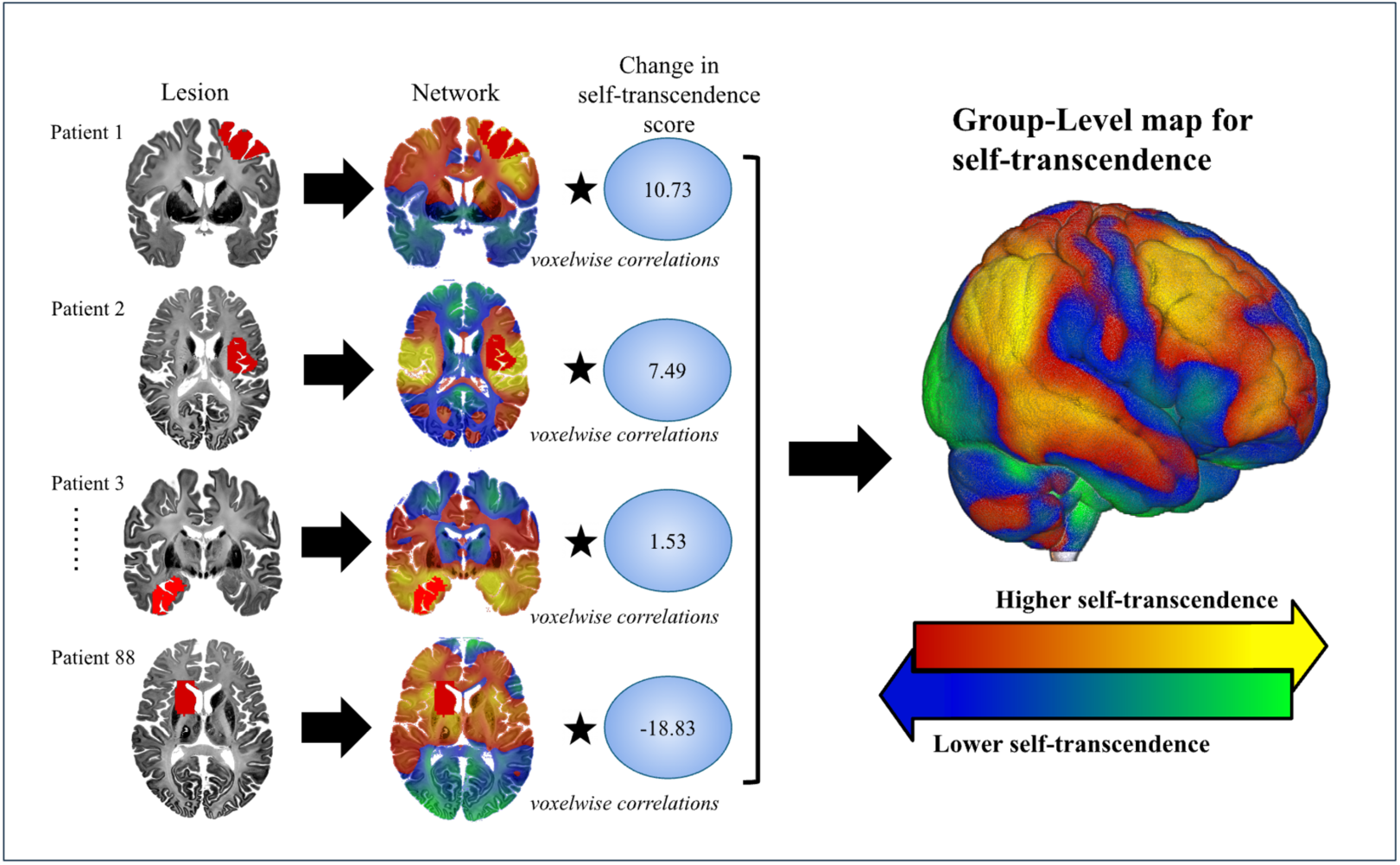
Data-driven method for identifying a lesion network for self-transcendence. (*Left*) The network of brain regions functionally connected to each lesion was computed using resting-state functional connectivity data from a large database of healthy volunteers (N = 1,000). Lesions and lesion networks are shown for 4 of the 88 patients in the Urgesi et al., 2010 dataset. Corresponding networks and behavioral scores for these patients are displayed. Positively-connected voxels are shown in warm colors, while negatively-connected voxels are shown in cool colors. Connections associated with self-transcendence were then identified (*Right).* Warm colors in the group-level map indicate that functional connectivity with lesions is more likely associated with higher scores for self-transcendence. Cool colors in the map indicate that functional connectivity with brain lesions is more likely associated with lower scores for self-transcendence.

To identify a distributed brain network whose functional connectivity with lesion sites corresponds to pre-to-post-operative changes in self-transcendence, we computed the cross-subject correlation, at each voxel, between lesion-seed connectivity and individual pre-to-post-operative ST change scores. Each voxel thus received a statistic reflecting the strength of association between connectivity at that location and ST change across the 88 patients. Permutation Analysis of Linear Models (PALM) (Winkler et al., 2014) was used to generate whole-brain t-maps in which positive functional connectivity with lesion sites corresponds to increases in self-transcendence, and negative functional connectivity corresponds to decreases (Fig. 2). Resulting t-statistic maps were visualized in MRIcroGL (v1.2) (Rorden, 2025).

### Comparison with meta-analytic fMRI maps for self-referential processing

To identify neuroanatomical commonalities and differences between the functional architecture of self-transcendence and self-reference, we obtained fMRI activation maps from an independent meta-analysis of self-referential processes (Hu et al., 2016) via BrainMap.org. We additionally examined an automated fMRI meta-analysis map derived from 166 studies tagged for the term ’self-referential’ on Neurosynth, a large-scale automated neuroimaging meta-analysis platform (Yarkoni et al., 2011).

### Internal consistency testing

To assess the stability of voxelwise associations, a leave-one-out (LOO) procedure was implemented. For each iteration, one subject was removed, voxelwise correlations were recomputed using the remaining subjects, and voxels passing P < 0.05 (uncorrected) were identified. This process yielded, for each voxel, a stability score reflecting the proportion of LOO folds in which the voxel passed the threshold. A heat map was generated by assigning each voxel its LOO stability fraction, producing a continuous image representing the consistency of voxelwise associations across iterations. Voxels with a stability fraction of 1.0 (that is, passing P < 0.05 in 100% of LOO folds) were identified as maximally stable, and their spatial distribution and peak locations were recorded.

### Cluster identification

Binarized lesion network overlap maps (MNI152, 2 mm) were restricted to the brain using an FSL MNI152 brain mask, which was resampled as needed and conservatively eroded by 2 mm. Clusters were identified using 3D connected-component labeling (26-connectivity). A minimum cluster size threshold of 10 contiguous voxels was applied.

For each cluster, a representative coordinate was defined as the deepest interior voxel, identified via a Euclidean distance transform (accounting for voxel dimensions) as the voxel maximally distant from the cluster boundary. Ties were resolved by selecting the voxel nearest the cluster centroid. Coordinates were converted to MNI space using the NIfTI affine. Cluster size (voxel count) and maximum interior distance (mm) were recorded.

### External validations

#### Validation using functional imaging meta-analysis of self-transcending emotions

We searched BrainMap.org for meta-analyses of the most consistently identified self-transcending emotions: compassion, gratitude, and awe (Stellar et al., 2017; Van Kleef & Lelieveld, 2022), using these terms along with "self-transcending emotion" as search terms. The search yielded one eligible meta-analysis, examining compassion (Kim et al., 2020), from which volumetric brain masks were obtained in NIfTI format. The largest contiguous set of voxels was located in the brainstem between the periaqueductal gray and the ventral tegmental area. To assess the specificity of any observed overlap to self-transcendent traits rather than TCI traits more broadly, all analyses were repeated using the TCI subscale of self-directedness (SD), a trait reflecting autonomy, self-determination, and goal-directed behavior that contrasts with the outward, boundary-dissolving orientation of self-transcendence (Cloninger, 1994; Cloninger et al., 1993). The SD lesion network was generated using the same procedure described above, substituting change in self-directedness scores as the behavioral covariate of interest.

### Validation using functional imaging meta-analysis of ketamine-induced neural activity

Ketamine, an NMDA receptor antagonist, reliably produces acute dissociative and self-transcendent phenomenology, including ego dissolution, depersonalization, and altered boundaries of self-experience (Hack et al., 2023; Ionescu et al., 2018; Marguilho et al., 2023). To further assess correspondence with pharmacologically-induced states of altered self-experience, we examined spatial correspondence between the ST network and coordinates derived from a meta-analysis of neuroimaging studies of ketamine administration (Ait Bentaleb et al., 2024). MNI coordinates from the meta-analysis were extracted and their spatial relationship with the unthresholded ST map was evaluated using the same approach applied to the compassion coordinates as described above. This analysis tested whether regions consistently modulated by a pharmacological agent known to alter self-experience correspond to regions whose lesion connectivity is associated with changes in trait self-transcendence.

### Validation via posterior cingulate cortex neuromodulation

To provide qualitative contextual alignment with prior neuromodulation work, we examined spatial correspondence between the ST network and a posterior cingulate cortex (PCC) locus previously targeted using low-intensity transcranial focused ultrasound (tFUS) to modulate DMN organization and self-related experiential orientation (Lord et al., 2024). A spherical region of interest (3.5 mm) centered on the reported PCC coordinate (−8, −56, 26) was constructed in MNI space and overlaid onto the unthresholded ST map. Given the single-coordinate nature of the stimulation site, spatial correspondence was assessed descriptively to determine whether a region previously shown to acutely modulate DMN organization and self-related experiential orientation aligns with the network whose lesion connectivity corresponds with trait-level variation in self-transcendence.

#### Network correspondence analysis

Spatial correspondence between the lesion-derived ST network and canonical cortical networks (Yeo et al., 2011) was quantified using the Network Correspondence Toolbox (NCT) (Kong et al., 2025). We first tested our a priori hypothesis that the ST network would show significant spatial correspondence with the DMN, followed by exploratory correspondence analysis across the full Yeo 7- and 17- network parcellations. For each canonical network, raw spatial overlap was quantified using Dice coefficients.

Statistical significance was assessed using surface-based spin tests that preserve spatial autocorrelation. The 17-network parcellation was additionally examined to enable identification of specific subnetworks within broader canonical systems. Multiple comparison correction was performed using Benjamini-Hochberg correction. Because spin test p-values reflect overlap relative to a network-specific null distribution, Dice coefficients and p-values need not rank-order identically across networks of differing sizes and spatial extents.

### Exploratory comparisons for self-transcendence network and lesions associated with other neurobehavioral conditions

To assess broader neurobehavioral relevance, the ST network was compared with 32 previously published lesion-derived, symptom-specific networks. These networks were associated with distinct neurobehavioral conditions and drawn from independent datasets totaling 923 lesion cases. For each condition, similarity with the ST network was quantified by computing its spatial correlation with each of the 923 lesions’ whole-brain functional connectivity maps. This was followed by an ANOVA across the full dataset and post-hoc one-sample t-tests within each subset of symptom-specific lesion networks.

## Results

### Deriving the network for self-transcendence

Lesion network mapping across 88 neurosurgical patients revealed a distributed, whole-brain pattern of functional connectivity associated with pre-to-post-operative changes in self-transcendence (Fig. 2, right; Fig. 3a). ST change scores were approximately normally distributed across the cohort (Supplementary Fig. 1). The resulting network encompassed the inferior parietal regions previously identified by voxel-lesion symptom mapping (Urgesi et al., 2010), while extending to a broader set of cortical and subcortical regions whose normative connectivity with lesion sites corresponded to individual differences in ST change.

**Figure 3:**
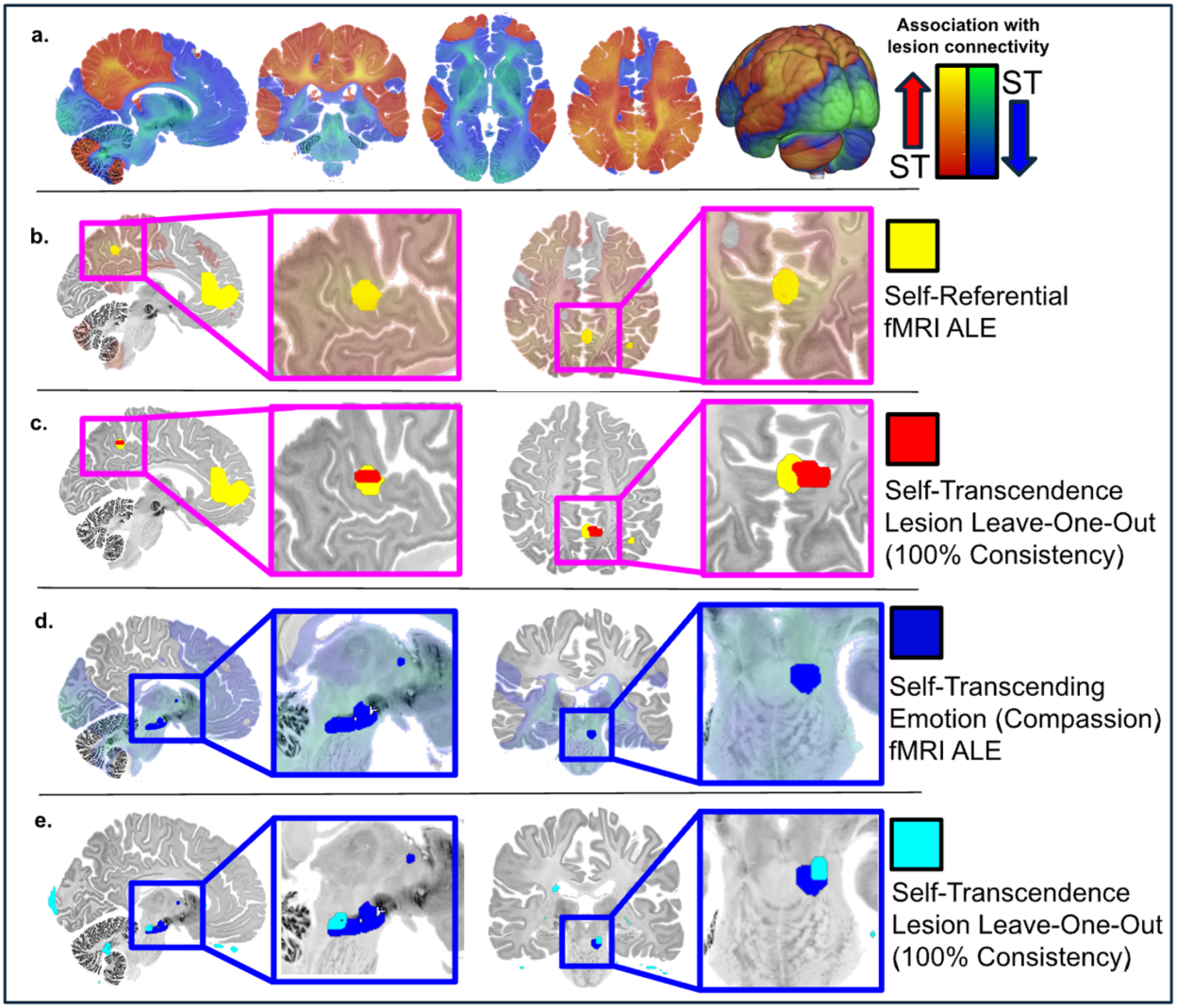
Convergent validity of the self-transcendence lesion network. (a) Whole-brain self-transcendence (ST) network derived from lesion network mapping. Warm colors indicate regions where functional connectivity with lesion sites corresponds with higher ST scores; cool colors indicate regions where connectivity corresponds with lower ST scores. Pink insets highlight regions of interest within the positive ST network; blue insets highlight regions within the negative ST network. (b) Spatial correspondence between the positive ST network and a meta-analytic map of self-referential processing (Hu et al., 2016). (c) Spatial correspondence between the self-referential processing meta-analytic map and a positive lesion leave-one-out consistency test (100% consistency). (d) Spatial correspondence between the negative ST network and a meta-analytic map of self-transcending emotions (compassion; Kim et al., 2020). (e) Spatial correspondence between the self-transcending emotion (compassion) meta-analytic map and a negative lesion leave-one-out consistency test (100% consistency).

### Comparison with meta-analytic maps for self-referential processing

Examination of meta-analytic maps of self-referential processing revealed spatial organization consistent with an anterior-posterior dissociation within the DMN, with regions associated with self-referential processing showing qualitative correspondence with the posterior-localized voxelwise effects within the lesion-derived ST network (Fig. 3b; Supplementary Fig. 2a,b).

### Internal consistency testing

The leave-one-out stability analysis demonstrated that a subset of DMN voxels consistently exhibited positive associations with ST change across folds. Although most voxels showed low stability, a small number displayed high stability fraction values, indicating reproducible associations despite leave-one-out perturbation of the dataset. Peak stable voxels were located in posterior medial cortical regions (precuneus and posterior cingulate vicinity), consistent with canonical DMN hubs. A small set of voxels passed the P < 0.05 threshold in all LOO iterations (stability fraction = 1.0), showed moderate effect sizes (mean r ≈ 0.22), and represented the most stable elements of the voxelwise association pattern (Fig. 3c).

The leave-one-out stability analysis additionally identified a subset of voxels consistently exhibiting negative associations with ST change across folds. Although most voxels showed low stability, a small number displayed high stability fraction values, indicating reproducible negative associations despite leave-one-out perturbation of the dataset.

### Cluster identification

Clusters meeting the ≥10 voxel threshold were identified in both positive and negative maps after restriction to a tightened intracerebral mask. Reported MNI coordinates correspond to the deepest interior voxel of each cluster, yielding anatomically valid, in-cluster representative locations (Table 1).

**Table 1:**
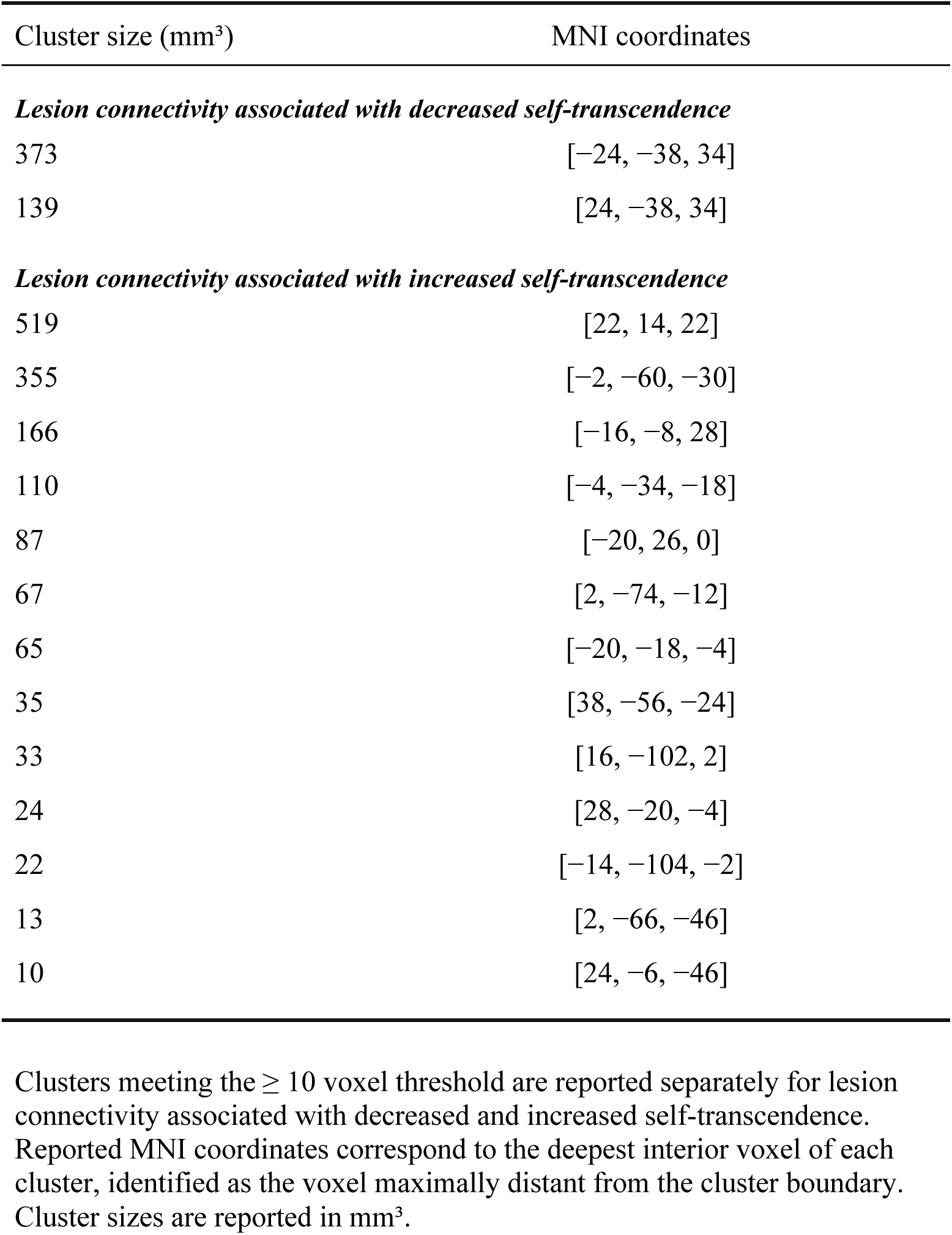
Peak cluster locations within the self-transcendence lesion network.

### External validations

*Validation using functional imaging meta-analysis of self-transcending emotions.* Coordinates derived from a meta-analysis of fMRI activation during compassion (Kim et al., 2020) showed a significant negative spatial association with the ST network (t = −6.370, P = 1.380 × 10⁻⁶, 95% CI [−1.100, −0.560]; Fig. 3d,e; Supplementary Fig. 2d). In contrast, compassion-related coordinates did not significantly overlap with the lesion-derived self-directedness network (t = −0.590, P = 0.560), supporting the specificity of this association to self-transcendence rather than TCI traits more broadly.

### Validation using functional imaging meta-analysis of ketamine-induced neural activity

Coordinates derived from a meta-analysis of neuroimaging studies of ketamine administration showed a spatial association with the ST network in the expected direction, though this did not reach statistical significance (t(4) = –2.486, P = 0.068). Visual inspection of the coordinate overlay revealed qualitative spatial correspondence between regions consistently modulated by ketamine and regions within the lesion-derived ST network (Supplementary Fig. 2e).

### Validation via posterior cingulate cortex neuromodulation

Consistent with the posterior midline localization of the ST network, the lesion-derived ST network encompassed the PCC locus previously targeted using low-intensity transcranial focused ultrasound to modulate DMN organization and self-related experiential orientation (Lord et al., 2024; Supplementary Fig. 2c).

### Network correspondence analysis

Consistent with our a priori hypothesis, the ST network showed significant spatial correspondence with the DMN (Dice coefficient = 0.393, P = 0.031), as quantified using the Network Correspondence Toolbox with surface-based spin tests preserving spatial autocorrelation (Fig. 4a; Table 2). Exploratory analysis of correspondence across the full Yeo 7- and 17- network parcellations revealed that the ST network showed significant correspondence with other canonical networks only at the 17- network level. These included Control B (Dice coefficient = 0.246, P = 0.001), Control C (Dice coefficient = 0.076, P = 0.003), Dorsal Attention B (Dice coefficient = 0.160, P = 0.007), and Somatomotor A (Dice coefficient = 0.232, P = 0.007) (Table 3; Fig. 4b). Lesion connectivity to these regions was associated with increased self-transcendence.

**Figure 4:**
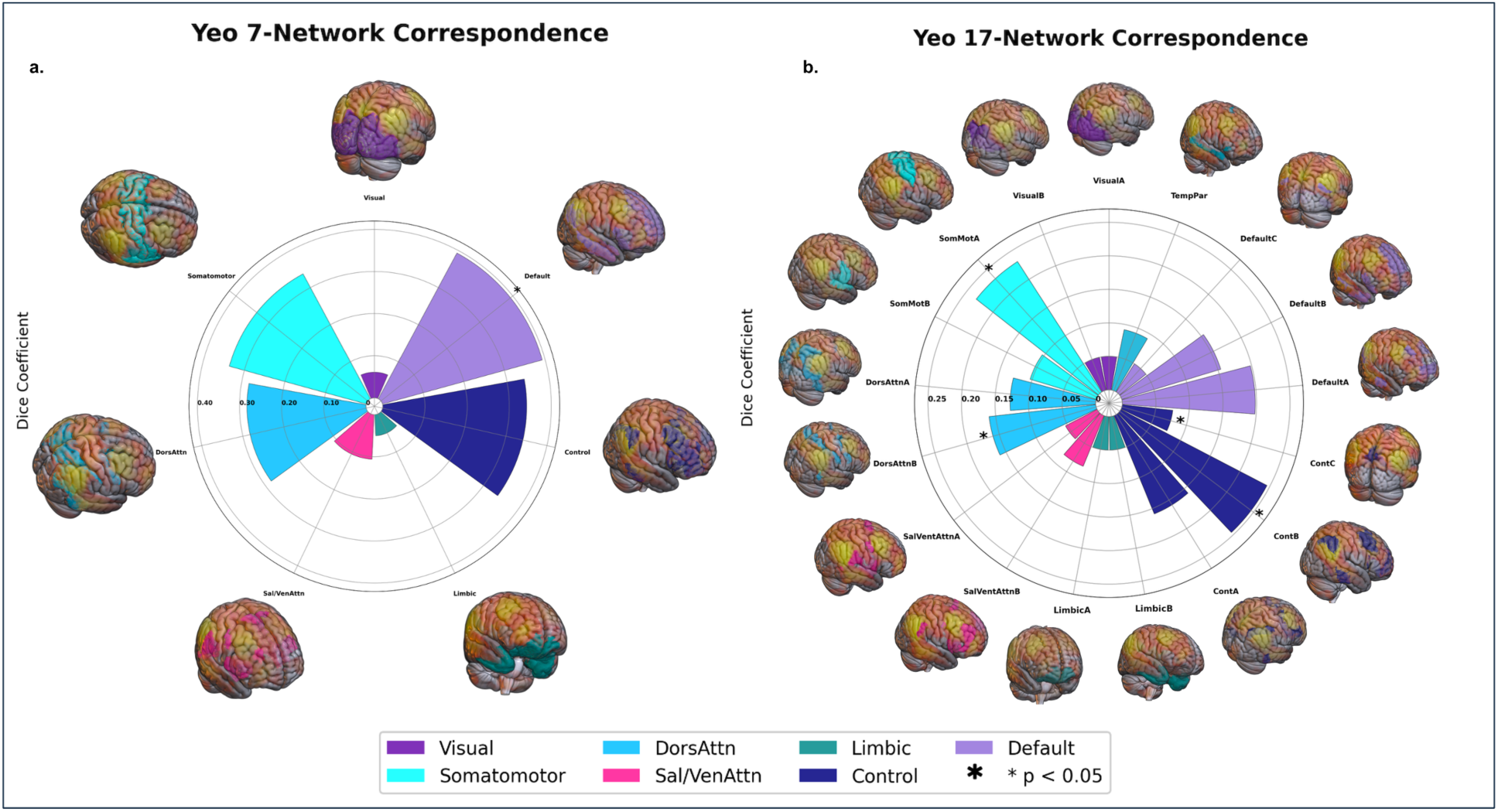
Network correspondence analysis of the self-transcendence lesion network with Yeo 7 and 17 canonical cortical networks. (a) Correspondence with the Yeo 7-network parcellation. Dice coefficients for each canonical network are displayed as radial bars. Consistent with our a priori hypothesis, significant correspondence was observed for the default mode network only (asterisk; *p_corrected_* < 0.05). The frontoparietal control network showed the second largest raw overlap but did not survive correction for multiple comparisons. (b) Correspondence with the Yeo 17-network parcellation. Exploratory analysis revealed significant correspondence (asterisk; *p_corrected_* < 0.05) for Control B, Control C, Dorsal Attention B, and Somatomotor A.

**Table 2:** Spatial correspondence between the self-transcendence lesion network and Yeo 7-network canonical cortical parcellations.

| Network | Dice Coefficient | P-value | BH Rank | BH Threshold | Significant |
| --- | --- | --- | --- | --- | --- |
| **Default** | 0.393 | 0.031 | primary hypothesis | NA | Yes |
| **Control** | 0.341 | 0.018 | 1 | 0.0083 | No |
| **DorsAttn** | 0.282 | 0.024 | 2 | 0.0167 | No |
| **Somatomotor** | 0.337 | 0.118 | 3 | 0.0250 | No |
| **Visual** | 0.061 | 0.875 | 4 | 0.0333 | No |
| **Sal/VenAttn** | 0.106 | 0.925 | 5 | 0.0417 | No |
| **Limbic** | 0.011 | 0.946 | 6 | 0.0500 | No |

Dice coefficients quantify raw spatial overlap between the self-transcendence (ST) lesion network and each of the seven canonical cortical networks. P-values were derived from surface-based spin tests that preserve spatial autocorrelation. Statistically significant correspondence ( $p_{corrected} < 0.05$ ) was observed for the default mode network only. The default mode network was tested as an a priori hypothesis and was therefore excluded from Benjamini-Hochberg correction, which was applied only to the remaining six networks. P-values reflect overlap relative to a network-specific null distribution and therefore need not rank-order with Dice coefficients across networks of differing sizes.

**Table 3:** Spatial correspondence between the self-transcendence lesion network and Yeo 17-network canonical cortical parcellations.

| Network | Dice Coefficient | P-value | BH Rank | BH Threshold | BH Significant |
| --- | --- | --- | --- | --- | --- |
| **VisualA** | 0.001 | 0.986 | 17 | 0.0500 | No |
| **VisualB** | 0.039 | 0.755 | 12 | 0.0353 | No |
| **SomatomotorA** | 0.232 | 0.007 | 3 | 0.0088 | Yes |
| **SomatomotorB** | 0.103 | 0.521 | 11 | 0.0324 | No |
| **DorsAttnA** | 0.127 | 0.353 | 8 | 0.0235 | No |
| **DorsAttnB** | 0.160 | 0.007 | 4 | 0.0118 | Yes |
| **Sal/VenAttnA** | 0.053 | 0.960 | 16 | 0.0471 | No |
| **Sal/VenAttnB** | 0.082 | 0.791 | 13 | 0.0382 | No |
| **LimbicA** | 0.001 | 0.937 | 15 | 0.0441 | No |
| **LimbicB** | 0.014 | 0.801 | 14 | 0.0412 | No |
| **ControlA** | 0.159 | 0.071 | 5 | 0.0147 | No |
| **ControlB** | 0.246 | 0.001 | 1 | 0.0029 | Yes |
| **ControlC** | 0.076 | 0.003 | 2 | 0.0059 | Yes |
| **DefaultA** | 0.198 | 0.158 | 6 | 0.0176 | No |
| **DefaultB** | 0.154 | 0.519 | 10 | 0.0294 | No |
| **DefaultC** | 0.039 | 0.517 | 9 | 0.0265 | No |
| **TempPar** | 0.092 | 0.284 | 7 | 0.0206 | No |

Dice coefficients quantify raw spatial overlap between the self-transcendence (ST) lesion network and each of the 17 canonical cortical networks. P-values were derived from surface-based spin tests that preserve spatial autocorrelation. Benjamini-Hochberg correction was applied across all 17 networks. P-values reflect overlap relative to a network-specific null distribution and therefore need not rank-order with Dice coefficients across networks of differing sizes.

### Exploratory comparisons for self-transcendence network and lesions associated with other neurobehavioral conditions

A one-way ANOVA across 32 previously published symptom-specific lesion networks revealed significant variation in spatial correlation with the ST network, F(30, 966) = 30.2, p = 4.6 × 10⁻^117^. Post-hoc testing identified loss of consciousness as having the strongest positive spatial correlation with the ST network (t = 2.28, P = 0.038). A positive trend was observed for alien limb syndrome (t = 1.738, P = 0.088) (Fig. 5).

**Figure 5:**
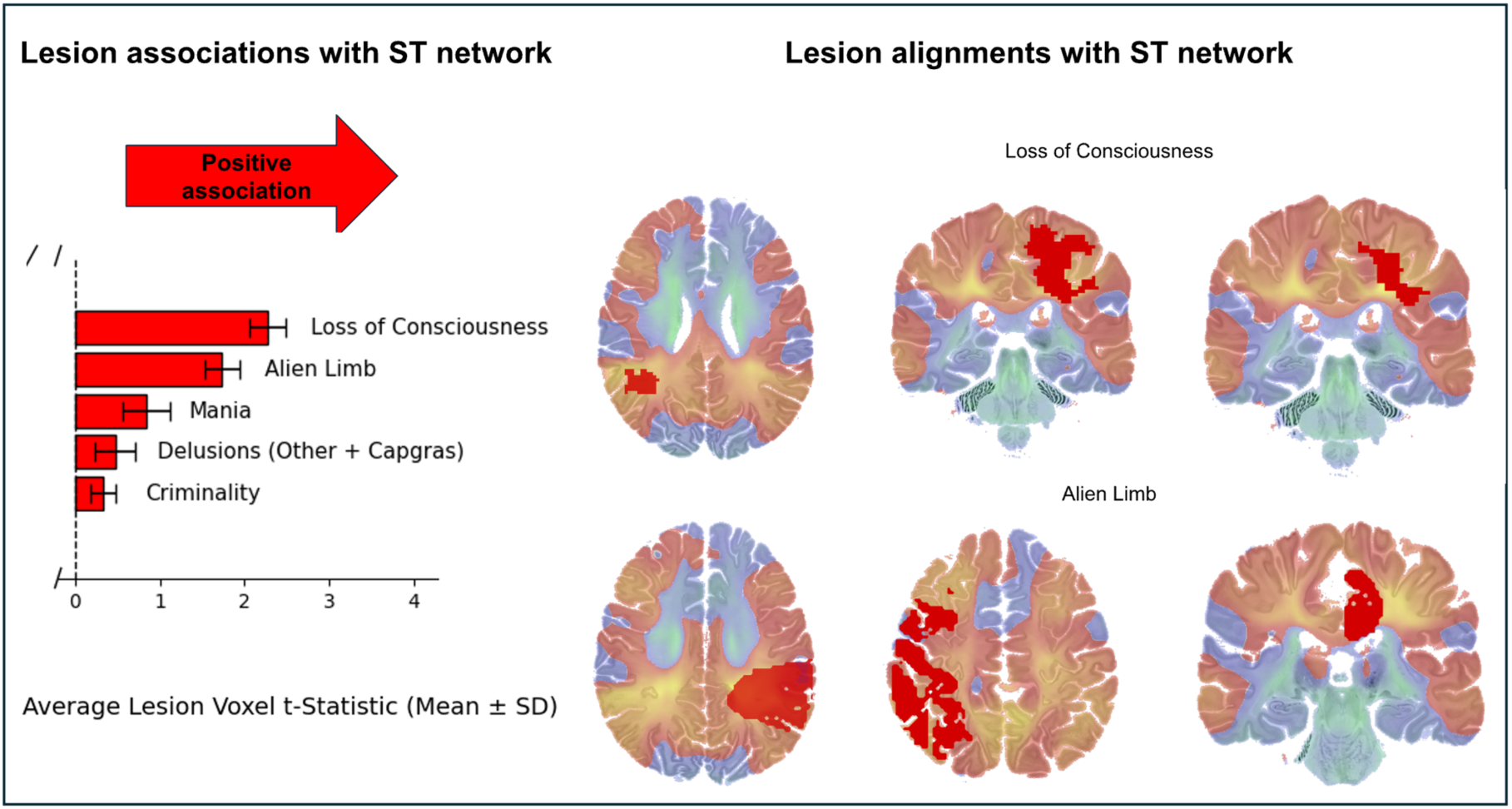
Exploratory comparisons of the self-transcendence network with 32 lesion-derived neurobehavioral networks. (*Left*) Bar graph displaying the average voxel t-statistics within lesion locations associated with neurobehavioral conditions. Error bars reflect standard error across lesion locations within each condition. The five conditions displayed represent the top five positive associations out of 32 lesion-derived neurobehavioral networks. The strongest positive association was observed with loss of consciousness, followed by alien limb syndrome. (*Right*) Spatial alignment between lesion locations and the self-transcendence (ST) network for the top two conditions. Top row: Lesions associated with loss of consciousness showed strong overlap with positive nodes of the ST network (warm colors), consistent with lesion locations associated with increased self-transcendence. Bottom row: Lesions associated with alien limb syndrome similarly overlapped with positive nodes of the ST network.

## Discussion

Using lesion network mapping in 88 neurosurgical patients, we identified a distributed brain network whose disruption increases self-transcendence. This network stably incorporates posterior cortical midline regions central to self-referential processing, spans multiple well-characterized cortical networks, and triangulates with independent functional, pharmacological, and neuromodulation evidence for the neural basis of self-transcendent experience. Together, this suggests an architecture in which posterior midline attenuation reduces self-referential processing, whereas engagement of key frontal midline and brainstem regions drives increased cognitive and affective expansion beyond self-reference alone.

Lesion connectivity associated with decreased self-transcendence implies that the intact function of connected regions normally supports self-transcendence. Conversely, lesion connectivity associated with increased self-transcendence points toward neuroanatomical regions where enhanced function would be expected to reduce self-transcendence. Lesions associated with decreased self-transcendence were stably connected with posterior cortical midline hubs of self-referential processing, implying their role in constraining self-transcendence, while sparing anterior self-referential hubs (Supplementary Fig. 2a,b). This anterior-posterior functional division of self-referential topography in our self-transcendence lesion network map is consistent with prior work demonstrating that these midline regions display different mechanistic roles in self-referential cognition (Northoff et al., 2006; Qin et al., 2012). Leave-one-out analyses confirmed this posterior connectivity pattern as the most stable association with increased self-transcendence following brain lesions (Fig. 3b,c). This stability pattern is consistent with prior work linking contemplative practices to reduced engagement of posterior midline self-referential systems (Bauer et al., 2019; Brewer et al., 2011; Garrison et al., 2015; Lord et al., 2026). Our result provides causal evidence that these changes are anatomically specific to posterior midline regions rather than reflecting uniform downregulation of self-referential processing.

Three independent lines of evidence converged on this network. First, coordinates from a meta-analysis of self-transcending emotion (compassion-related fMRI activation) aligned with the ST network in the direction predicted by lesion network mapping logic (Fig. 3d; Supplementary Fig. 2d). Specifically, the most stable associations with post-lesion decreases in self-transcendence localized to brainstem regions recruited by compassion (Fig. 3d,e). In other words, the brainstem regions that are recruited by self-transcending emotion are also the ones most stably associated with reduced self-transcendence when they are connected to lesions in our dataset. Second, coordinates from neuroimaging meta-analysis of ketamine administration showed a spatial association trending with the ST network in the expected direction (Supplementary Fig. 2e). The failure to reach significance likely reflects the limited number of available coordinate clusters rather than absence of true correspondence. Third, the ST network encompassed a PCC locus previously targeted using transcranial focused ultrasound to modulate DMN organization and the sense of self (Lord et al., 2024) (Supplementary Fig. 2c).

Consistent with our a priori hypothesis, the ST network showed significant spatial correspondence with the DMN (Fig. 4a). There was an association with frontoparietal control networks identified during exploratory analysis, though it did not survive multiple comparison corrections. The involvement of these two networks is consistent with the original inferior parietal lesion locus identified by Urgesi et al. (2010), which falls at the intersection of the default mode and frontoparietal control networks (Supplementary Fig. 3). The 17-network characterization demonstrated significant associations between the ST network and a wider range of functional network sub-units (Fig. 4b). Notably, while the ST network showed significant correspondence with the DMN at the 7-network level, no individual DMN subnetwork survived correction at the 17-network level, suggesting that the ST network’s relationship with the DMN is topographically distributed across its subcomponents.

In contrast, frontoparietal correspondence strengthened with division into subnetworks, suggesting that the ST network’s correspondence with the frontoparietal system is topographically concentrated within specific subnetworks. These results correspond to observations that the frontoparietal control network serves as an intermediary between the internally-oriented DMN functions and externally-oriented dorsal attention networks (Cole et al., 2013; Marek & Dosenbach, 2018; Vincent et al., 2008), and provides support for theoretical models that implicate both self-referential and regulatory systems in self-transcendent phenomena (Newberg & Yaden, 2018; Vago & Silbersweig, 2012).

Exploratory comparisons with 32 previously published symptom-specific lesion networks revealed that the ST network was most similar to lesion networks associated with loss of consciousness, with a trend-level association with alien limb syndrome (Fig. 5). Both conditions involve disruptions in the coherent binding of self-experience (Z. Huang et al., 2014; Olgiati et al., 2017). The network similarity with loss of consciousness is consistent with research operationalizing the cessation of conscious self-experience as a meditative endpoint (Laukkonen et al., 2023; Shinozuka et al., 2025).

Several limitations warrant consideration. First, self-transcendence was assessed using a trait-based self-report measure, and the relationship with state-based or experimentally induced forms merits considerable future research. Second, our exploration of self-transcendent emotions was limited to one out of the three widely recognized self-transcending emotions, as no fMRI meta-analyses have been published on awe or gratitude. Third, lack of a sufficiently powered neurosurgical independent dataset with corresponding longitudinal changes assessed in the TCI prevents a direct network replication. We mitigate this through the various converging lines of evidence reported in the manuscript. Fourth, the study relies on pre- to post-operative change in TCI self-transcendence scores as the behavioral variable of interest. Because TCI self-transcendence is typically considered a relatively stable personality trait, observed post-surgical changes may partially reflect non-specific factors associated with the surgical experience itself, including acute stress, mood fluctuation, anesthesia effects, or post-operative medication, rather than lesion-specific effects on self-transcendence per se. Future studies should incorporate control measures of surgery-related psychological change and, where feasible, longitudinal follow-up assessments to distinguish trait-level reorganization from transient perioperative effects.

Together, these findings establish a network architecture for self-transcendence. Triangulation of evidence across lesion-derived connectivity, meta-analytic activation maps, and neuromodulation targets supports our model of self-transcendence as a phenomenon involving constraint by posterior midline cortical hubs and facilitation by brainstem and anterior midline cortical hubs. This model offers a unified framework that both accounts for previously disparate findings and generates novel, testable predictions about how the brain gives rise to self-transcendence.

## Supplementary Figures

**Supplementary Figure 1:**
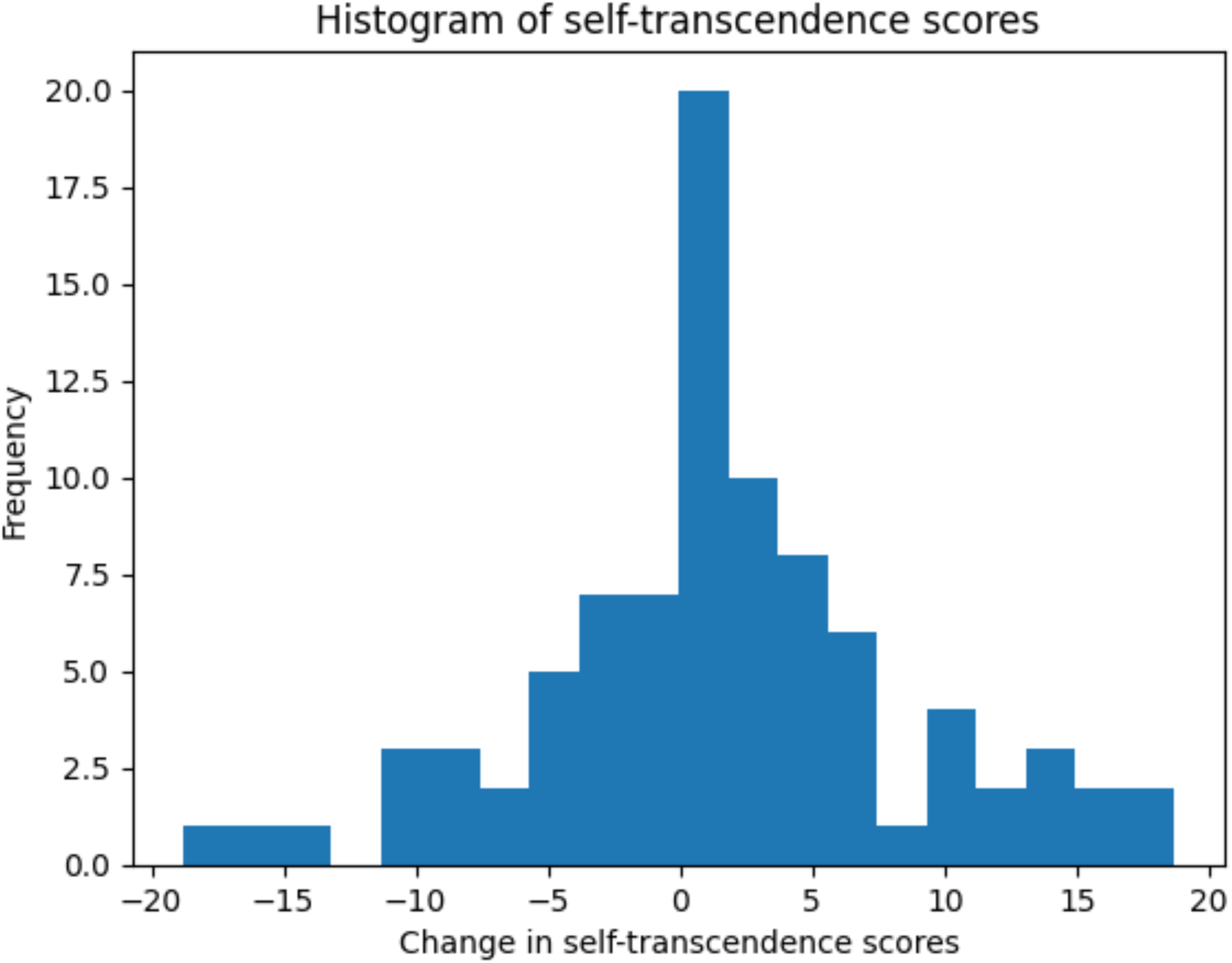
Distribution of pre- to post-operative change in self-transcendence scores. Histogram showing a normal distribution of individual changes in Temperament and Character Inventory self-transcendence scores across the lesion cohort post- vs pre- surgery.

**Supplementary Figure 2:**
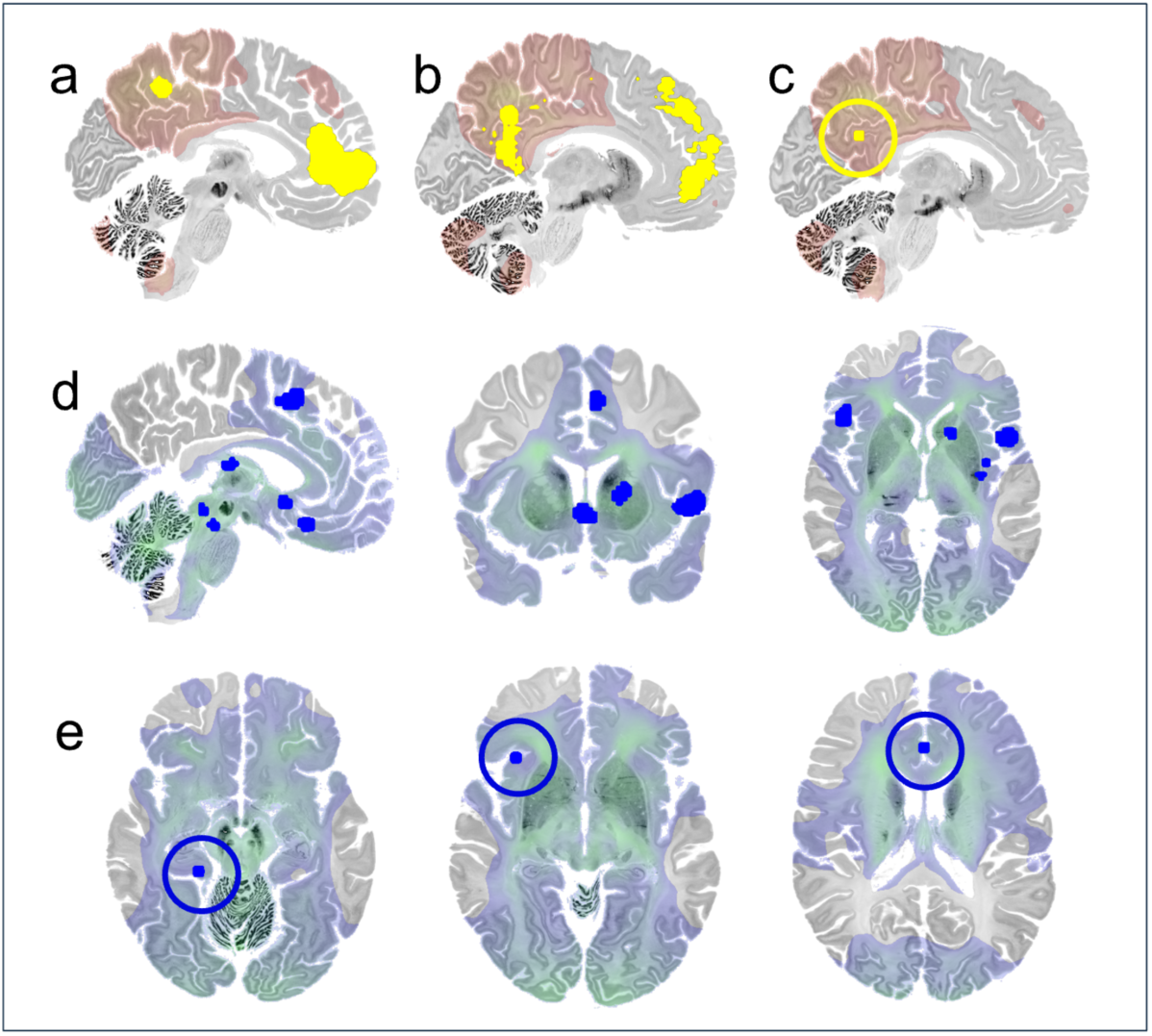
Convergent validity of the self-transcendence lesion mapping across independent sources. (a) Spatial correspondence between positive zones of the ST network and a meta-analytic map of self-referential processing (Hu et al., 2016), shown in sagittal view. Yellow regions indicate overlap between the ST network and self-referential processing activation coordinates. (b) Spatial correspondence between positive zones of the ST network and a Neurosynth-derived meta-analytic map of self-referential processing, shown in sagittal view. (c) Qualitative spatial correspondence between the positive ST network and the posterior cingulate cortex (PCC) neuromodulation stimulation site associated with altered experience in self representation (Lord et al., 2024), indicated by a yellow circle. (d) Spatial correspondence between negative zones of the ST network and a meta-analytic map of compassion-related activation (Kim et al., 2020), shown in axial views. Blue regions indicate overlap between the ST network and compassion activation coordinates. (e) Spatial correspondence between negative zones of the ST network and coordinates derived from ketamine studies, shown in axial views. Blue circles indicate regions of spatial convergence.

**Supplementary Figure 3:**
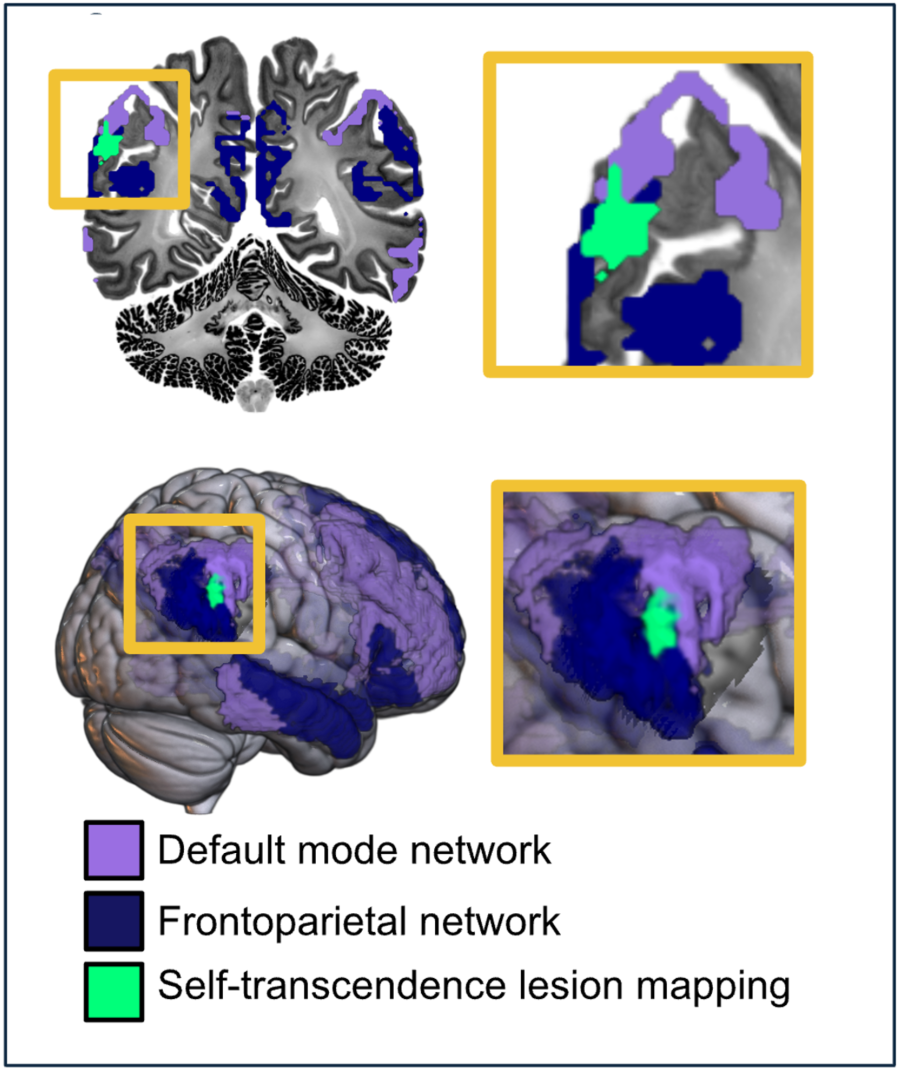
Convergence of self-transcendence lesion mapping, default mode network, and frontoparietal networks. Overlap between the self-transcendence lesion mapping spot identified by Urgesi et al. (2010) (green) and the Yeo-7 default mode network (light purple) and frontoparietal control network (dark blue) shown in coronal slice (top row) and lateral surface (bottom row) views. Orange insets provide zoomed views of the region of convergence. All three maps converge on inferior parietal cortex, demonstrating that the original lesion locus associated with self-transcendence falls at the intersection of the default mode and frontoparietal control networks.

